# Proteomic Analysis of NRROS Interactome Reveals the Presence of Chaperones and Mediators of the ERAD Pathway

**DOI:** 10.1101/326512

**Authors:** Rajkumar Noubade, Qui Phung, Wilson Phung, Erik Verschueren, Laura Lau, Hiroyasu Konno, Jennie Lill, Wenjun Ouyang

## Abstract

Negative regulator of reactive oxygen species (NRROS, previously called LRRC33) is a leucine-rich repeat (LRR) domain containing, ER-resident transmembrane protein expressed primarily in lymphoid organs, especially in myeloid cells. We have previously demonstrated that NRROS regulates reactive oxygen species production by phagocytic cells by mediating degradation of NOX2 (gp91^phox^), a component of NOX2 complex responsible for the oxidative burst in these cells. Since LRR is the only functional domain in NRROS, it is likely to interact with other proteins for its biological functions. Here, by performing immunoprecipitation of NRROS and mass spectrometric analysis, we describe the NRROS interactome in macrophages and demonstrate that NRROS interacts with molecular chaperones/co-chaperones and mediators of the endoplasmic reticulum associated degradation (ERAD) pathway such as calnexin, suggesting a broader role for NRROS in protein biosynthesis and the ER quality control machinery.

## Introduction

Negative regulator of reactive oxygen species (NRROS, previously known as Leucine Rich Repeat Containing protein 33 (LRRC33)) is a leucine-rich repeat (LRR) domain containing transmembrane protein. NRROS protein contains 22 putative LRR domains, a transmembrane domain, and a short cytoplasmic domain^1^. The LRR domain is present in many proteins, especially in proteins associated with innate immunity^2^. The LRR is a 20-29-aa motif and contains a conserved 11-aa sequence, LxxLxLxxN/CxL (where x is any amino acid)^3^. Most of the LRR domains occur in tandem arrays of two to more than a dozen chains^4^. Multiple LRRs can form a non-globular, horseshoe-shaped structure, wherein parallel beta-sheets line the inner circumference of the horseshoe and alpha helices. The function of many LRR domains is to provide a structural framework for protein-protein interactions^5^.

We have previously reported that NRROS is expressed primarily in lymphoid organs, especially in myeloid cells including macrophage subsets and neutrophils. NRROS is preferentially located in the endoplasmic-reticulum (ER)^1^. Functionally, we demonstrated that NRROS regulates reactive oxygen species (ROS) production by myeloid cells and that macrophages and neutrophils deficient of NRROS produce significantly more ROS in response to various stimuli. Although the increased ROS improves host defense in mice lacking NRROS, it exacerbated disease development in an experimental autoimmune encephalomyelitis (EAE) model. Recently, NRROS is also identified to exert essential roles in controlling the development of microglia during early embryonic stage^6^. The brains from NRROS-deficient mice are devoid of normal microglia, and instead are populated with perivascular macrophages. Although these mice develop normally till week 6–8, they display neurological abnormalities and succumb prematurely before 6 months of age. Mechanistically, at the molecular level, we showed that NRROS regulates ROS by regulating protein stability of a member of the NADPH oxidase 2 (NOX2) complex, a multi-protein complex responsible for the oxidative burst in phagocytic cells. The NOX2 complex is composed of two membrane-bound subunits, NOX2 (gp91^phox^) and p22^phox^, and four cytosolic subunits, p40^phox^, p47^phox^, p67^phox^ and Rac (Rac1 or Rac2).^7–9^ NOX2 and p22^phox^ form a heterodimer known as flavocytochrome b_558_ in the ER, and the hetero-dimerization is essential for the stability of individual monomers which otherwise get degraded.^10–12^ We showed that NRROS binds to NOX2 monomer and increases its degradation through an ER Associated Degradations (ERAD)-mediated process^1^.

Given the role of LRR motifs in protein-protein interactions and the observation that NRROS regulates NOX2 post-transcriptionally, we wished to understand the NRROS interactome to gain further insights into it functionalities. Here, through affinity purification of NRROS coupled with mass spectrometric analysis, we demonstrate that NRROS interacts with molecular chaperone/cochaperones and mediators of ER Associated Degradations (ERAD) pathway such as calnexin and speculate NRROS to be a molecular chaperone in the biosynthesis of multi subunit proteins such as NOX2 complex, either by itself or in association with calnexin.

## Material and Methods

### Plasmids

Sequence coding murine NRROS protein was amplified by PCR using mouse splenic cDNA as template, and cloned into expression vector pCDNA3 or retroviral vector pMSCV as described before.^1^ Similarly, HA tag was inserted at the c-terminus of mouse TGFβ1 that was amplified from template obtained from Origene and cloned into pCDNA3.1(+) vector using Infusion kit (Clonetech). NRROS^LAP^ constructs were generated using strategy as described.^13–14^ Specifically, NRROS cDNA was first cloned into the Gateway vector pDONR221 (Invitrogen). Then expression constructs were generated by Gateway cloning into pG-LAP5 (pEF5-Speptide-X-PrecissionS-EGFP). For retroviral constructs, the C-terminal GFP tagged NRROS^LAP^ was subcloned into pBABE Puro. All constructs were verified by sequencing and expressed and purified using standard methods.

### Generation of stable cell lines

RAW cells expressing NRROS^Flag^ or NRROS^LRR-Flag^ or NRROS^LAP^ were generated as described before.^1^ Briefly, Phoenix E cells were transfected using calcium phosphate method with one of the following constructs: NRROS^nFlag^-pMSCV, NRROS^LRR-Flag^-pMSCV, GFP-pMSCV control vector, NRROS^LAP^ pBABE or pBABE control plasmid. After 48 h, the retroviral supernatants were harvested, filtered through 0.45 μm filters, supplemented with polybrene (10 μg/ml) and added to RAW cells that were plated in a six well plate previous night. Plates were centrifuged at 1,200 × g at 32°C for 120 min. The viral supernatant was replaced by complete DMEM after an additional 2 h. Transduced RAW cells were sorted (FACSAria, BD Biosciences) based on GFP expression to generate polyclonal stable cells lines. For generation of NRROS^LAP^-low expressers, single cells were sorted into 96 well plates to generate clonal stable cell lines. Then colonies that grew were screened for GFP transcript expression using cells-to-CT method (Thermo Fisher). Subsequently, 48 positive clones were grown in 6-well plates and screened for NRROS^LAP^ expression by Western blot analysis using anti-NRROS antibody.^1^ Clone 12 was selected for further studies, based on similar expression level NRROS^LAP^ to that of endogenous NRROS as quantified by densitometry.

### LAP Tandem affinity purification

Stable RAW 264.7 cells expressing NRROS^LAP^ or GFP^LAP^ were pelleted (~2 ml packed cell volume) and pellets were snap frozen. LAP purification with anti-GFP antibody (kind gift from Dr. Peter Jackson’s lab), PreScission protease cleavage, and subsequent Protein S agarose affinity purification were performed according to the protocol described.^15 14^ Briefly, lysates generated from the frozen cell pellets were subjected to immunoprecipitation using anti-GFP antibody crosslinked to Affi-Prep Protein A beads (Bio-Rad). The washed beads were treated overnight with PrecissionS and digests were eluted. The eluted sample was further incubated with S protein agarose for 3 hours. Then washed agarose beads were subjected to urea elution and eluted material was used for downstream analyses including mass spectrometry analysis.

### Transfection of cells

293T cells grown in 100 mm plates were transfected with the indicated plasmid using Lipofectamine 2000 (Life Technologies) according to manufacturer’s instructions. Briefly, 10 μg total DNA was diluted in 500 μl of opti-MEM and 25 μl of Lipofectamine 2000 was diluted in another 500 μl of opti-MEM. The two were combined, incubated for 20 minutes and added drop-wise to cells grown on 100 mm plates at 70–80% confluency. After 24 h, cells were harvested by trypsinization and lysed in RIPA buffer (50 mM Tris pH 7.4, 150 mM NaCl, 2 mM EDTA, 1% NP-40, 1% SDS) and subjected to Western blot analysis or were used for immunoprecipitation with the indicated antibody.

### Anti-Flag Immunoprecipitation, Silver staining and Western blot analysis

Cell lysates were generated by lysing cells in in lysis buffer (50mM Tri-HCl, pH7.4, 150mM NaCl, 1% Triton X-100, 1mM EDTA Tris, EDTA,) containing protease and phosphatase inhibitor cocktail (Halt protease and phosphatase inhibitor cocktail from Pierce). For anti-FLAG immunoprecipitation, lysates were incubated with EZview Red ANTI-FLAG M2 affinity gel (Sigma) at 4°C for 4 h and eluted with 3X FLAG peptide (Sigma) according to manufacturer’s instructions. About 10% of the eluted material was run on 4–20% Bis-Tris gel (Invitrogen) and proteins were visualized by silver staining using SilverQuest silver staining kit (Life Technologies), according to manufacturer’s instructions. For western blot analysis, equal amounts of proteins, measured using BCA assay (Pierce), were separated by SDS-PAGE on 4–12% Bis-Tris gels (Life Technologies). and transferred onto nitrocellulose membranes using iBlot apparatus (Invitrogen). Membranes were blocked for 1 h at room temperature with 5% milk in TBST and probed with indicated antibody in 5% milk overnight at 4°C. The primary antibodies used included NOX2 (gp91^phox^) (clone 54.1,), p22^phox^ (clone FL-195), GFP (clone B-2) from Santa Cruz Biotechnology, anti-HA (clone C29F4), GAPDH-horseradish peroxidase (HRP) (clone 14C10), HSP90 (clone C45G5), HSP70 (clone D69), CHIP (clone C3B6) from Cell Signaling Technology, calnexin (rabbit polyclonal) from Stressgen Biotechnonlogies, actin (rabbit polyclonal) from Sigma-Aldrich. Anti-NRROS monoclonal antibody was generated in-house^1^. Secondary antibodies include HRP-conjugated anti-rabbit, anti-mouse (Cell Signaling Technology) and anti-Armenian hamster (Jackson Immunoresearch).

### Mass spectrometric analysis

Anti-Flag or LAP-tagged immunoprecipitates from RAW264.7 expressing tagged NRROS or GFP as a control were reduced with 10mM dithioreitol (DTT) for 1 h at 37^0^ C, and alkylated with N-isopropyliodoacetamide at room temperature in the dark for 30 min. Samples were then loaded onto a 4–12% Bis-Tris gel (Life Technologies). Entire gel lanes were excised and divided from top to bottom into 11–20 bands. The gel bands were de-stained with 50:50 methanol:50mM ammonium bicarbonate in water then digested at 37°C overnight with 0.02 mg/mL trypsin (Promega) in 50mM ammonium bicarbonate. The digested samples were injected onto a 100 μm inner diameter capillary column (NanoAcquityUPLCcolumn, 100 μm x100mm, 1.7 μm, BEH130 C18, Waters Corp) and separated by capillary reverse phase chromatography on a NanoAcquity UPLC system (Waters Corp). Samples were eluted of the column with a gradient of 2–90% buffer B (in which buffer A is 0.1% formic acid, 2% acetonitrile and 98% water, and buffer B is 0.1%formic acid, 2% water and 98% acetonitrile) at 1 μl/min with a total analysis time of 45 min. Peptides were eluted directly into an LTQ-Orbitrap XL mass spectrometer (ThermoFisher) and ionized using an ADVANCE source (Michrom-Bruker) at a spray voltage of 1.4 kV. Mass spectral data were acquired using a method comprising of one full mass spectrometry scan (375-1,600 m/z) in the Orbitrap at 60,000 resolution M/ΔM at m/z 400 followed by collision-induced of the top 8 most abundant ions detected in the full mass spectrometry scan in a cycle repeated throughout the liquid chromatography gradient in the linear ion trap. Tandem mass spectral results were submitted for database searching using the Mascot search algorithm ver 2.4.1 (Matrix Sciences) against a concatenated target-decoy database (Uniprot ver 2011_12) consisting of murine proteins and common laboratory contaminants such as trypsin. The data was searched with tryptic specificity, allowing two mis-cleavages, variable modifications of cysteine NIPCAM (+99.0684 daltons (Da)), methionine oxidation (+15.9949 Da), lysine ubiquitylation (+114.0429 Da), serine, threoninin, tyrosine phosphorylation (+79.9663 Da) 50ppm precursor ion mass, and 0.8Da fragment ion mass tolerance specified. Peptide spectral matches were filtered using a linear discriminant algorithm to an estimated peptide false discovery rate (FDR) of 5% followed by protein-level FDR of 2%. All Peptide Spectral Matches (PSMs) per protein were summed per sample across all fractions from the GelC-MS experiment. The Statistical Analysis of INTeractome (SAINT) algorithm (SAINTExpress-spc v.3.6.1)^16^ was run with default settings comparing the sum of PSMs for all identified proteins enriched with each antibody separately (target) to the combined pool of control purifications. Interactions with a SAINT Bayesian False Discovery Rate (BFDR) < 0.05 were marked as significant.

## Results

To gain insights into the NRROS interactome, we focused our efforts on RAW 264.7, a macrophage-like murine cell line, since NRROS is highly expressed in myeloid cells such as macrophages and neutrophils.^1^ To aid the immunoprecipitation, we created epitope-tagged NRROS to be used as bait and, utilizing retroviral transduction, generated two independent stable cell lines overexpressing NRROS: one with Flag-tag at the N-terminus of the protein (NRROS^nFlag^) and another with Flag-tag internal to the protein (NRROS^LRR-Flag^). The expression of NRROS^nFlag^ and NRROS^LRR-Flag^ in those cell lines was confirmed by western blot analysis (**Figure 1A**). No difference in NRROS expression or localization between the two cells lines was observed (data not shown). Whole cell lysates from the stable cells were subjected to affinity purification (AP) using anti-Flag antibody, followed by mass spectrometry (AP-MS) analysis to identify NRROS-interacting proteins. Consistent with expectations, NRROS as bait was immunoprecipitated from cells expressing tagged-NRROS, with unique peptide coverage of 43–45% (**Table S-1**) but not from control cells. We considered NRROS^nFlag^ and NRROS^LRR-Flag^ immunoprecipitations as replicates (n=2) and analyzed AP-MS data using the Statistical Analysis of INTeractome (SAINT) algorithm SAINTExpress.^16^ The logOddsScore for all significant proteins (FDR < .05) revealed several chaperones/co-chaperones and mediators of the ERAD pathway in the NRROS interactome. The proteins that interacted with NRROS included calnexin, UDP-glucose/glycoprotein glucosyl transferase (UGGT), Hypoxia upregulated 1 (HYOU1), T-complex proteins, and Dnajb1 (**Figure 1B and 1C)**. Among the top hits with FDR< 0.05, the SAINTExpress analysis showed calnexin to be a high-confidence interactor (SAINT score = 1) with 41% coverage in the NRROS^nFlag^ sample (267 total and 33 unique peptides) and 48% coverage in the NRROS^LRR-Flag^ sample (173 total and 25 unique peptides) (**Table S-1**). Additional proteins that interacted with NRROS included molecular chaperones such as Binding Immunoglobin protein (Bip)/Glucose-regulated protein 78(GRP78)/HSPA5 and protein disulfide isomerase (PDI) family members (**Table S-2**). Interestingly, a notable interactor that does not belong to the ERAD and protein folding pathways was Tgfβ1. However, while there were significantly higher number of unique peptides for the chaperone/co-chaperone molecules in NRROS-expressing cells, a few peptides for some of the same proteins, including calnexin, were observed in the control cells, leading to concerns about potential false positive hits. In order to reduce the background and non-specific interactions, we next utilized the Localization and Affinity Purification (LAP)-tag technology.^13 14^ The LAP tag consists of GFP, which aids in cell isolation and immunoprecipitation using anti-GFP antibody, PreScission protease cleavage site for native protein elution and S-peptide epitope for second affinity purification (**Figure 2A**). We constructed C-terminus LAP-tagged NRROS (NRROS^LAP^) in a retroviral vector and generated stable RAW 264.7 cell lines overexpressing NRROS^LAP^. Upon tandem affinity purification, first with anti-GFP antibody and then with S-agarose beads, we confirmed efficient immunoprecipitation by visualization of the enriched bait protein using silver stain analysis (**Figure S-1**). The protein-protein interactions were identified by mass spectrometry analysis, which revealed good coverage of the bait protein NRROS^LAP^ (53% protein coverage, with 688 total peptides and 37 unique peptides). Similar to the results obtained with NRROS^Flag^, NRROS^LAP^-interactome consisted of ER/mitochondrial chaperone/co-chaperone proteins and mediators of the ERAD pathway (**Table S-3**). The NRROS^LAP^-associated proteins included calnexin, Bip, UGGT1, Hyou1, T-complex proteins, Dnajb1, EDEM1 and members of Hsp90 and Hsp60 family. The interactome also included TGFβ1. In the GFP^LAP^-expressing control cells, unlike the Flag-epitope tagged approach, we observed no or significantly reduced peptides for the interacting proteins. For our representative, high-confidence interactor calnexin, we observed 49% coverage of the protein (212 total peptides with 42 unique peptides) (**Table S-4**). The similar NRROS-interactome obtained from two completely independent approaches point towards ER as the primary cellular compartment of NRROS functions. However, since NRROS is localized to the ER membrane, ^1^ it was possible that some of the proteins identified in the interactome were artifacts of ectopic NRROS overexpression. Therefore, we next generated cell lines that would express NRROS^LAP^ at levels close to that of endogenous protein. Since NRROS^LAP^ has higher molecular weight than endogenous NRROS due to the presence of GFP tag, an anti-NRROS antibody was used to detect and differentiate between the endogenous NRROS and exogenous NRROS^LAP^. Single cells were sorted, post transduction with NRROS^LAP^, based on GFP expression and large number of colonies (~200) were screened by assaying GFP transcripts (data not shown). Next, a selected number (~50) of positive colonies were expanded and screened for the presence of NRROS^LAP^ by western blot using anti-NRROS antibody (**Figure 2b**). Clone 12 was selected (**Figure 2b**) for further analysis based on comparable expression of ectopic NRROS^LAP^ and endogenous NRROS as determined by densitometry. Lysates from clone12 cells were subjected to sequential immunoprecipitation of NRROS^LAP^ with anti-GFP antibody and S-peptide agarose beads, efficient immunoprecipitation was confirmed by western blot (**Figure 2c**) and visualized bait protein enrichment by silver staining (**Figure 2d**). Mass spectrometric analysis showed good coverage of the bait protein NRROS (54% coverage with 382 total and 41 unique peptides) (**Table S-5**). The NRROS interactome, consistent with the previous results, was comprised of chaperones/co-chaperones proteins and mediators of ERAD pathway including calnexin, Bip, T-complex proteins and members of Hsp90 and Hsp60 family (**Table S-6**). The AP-MS analysis yielded 41% coverage of calnexin (33 total and 24 unique peptides) (**Table S-5**). Overall, the number of total peptides for NRROS^LAP^ was lower in this experiment compared to the other LAP-tag experiment, possibly pointing towards lower NRROS^LAP^ expression and recovery (**Table S-4** vs **Table S-5**). Additionally, while the NRROS interactome was consistent with the other two AP-MS analyses, the overall number of peptides was also generally low for all the interactors (**Figure 3A**). However, since all three approaches yielded similar NRROS-interactome, we compared the overlap between datasets. Comparison of NRROS^Flag^ dataset with the two NRROS^LAP^ AP-MS analyses yielded 19 proteins that were identified in NRROS^Flag^ dataset and in one or both NRROS^LAP^ datasets (**Table 1**). Further, there were 8 proteins that were common between all three datasets (**Figure 3A** and **Table 1**). Gene Ontology (GO) term analysis of the interactome indicated unfolded protein binding as the most probable molecular function (**Figure 3B**). Recently, interactomes of a large number of human proteins have been described in the BioPlex 2.0 database.^17^ This database includes NRROS (LRRC33) as one of the bait proteins in their AP-MS analysis. Integrating our data with the Bioplex 2.0 dataset yielded a 7-protein interactome network for NRROS (**Figure 3C**). Taken together, these data demonstrate that NRROS interactome includes several molecular chaperones and co-chaperones in the ER.

**Table 1.** Overlap between significant interactions in NRROS^Flag^ (n=2, FDR < .05 in SAINT analysis), NRROS^LAP^ over-expression (all proteins) and NRROS^LAP^ clone 12 (expression at endogenous level) (all proteins). The highlighted proteins are common to all three datasets.

| Interactor | NRROS<sup>Flag</sup> | NRROS<sup>LAP</sup> | NRROS<sup>LAP</sup> Clone 12 |
| --- | --- | --- | --- |
| PSA3 | 4.5 | NA | 1 |
| TCPE | 6 | 9 | 2 |
| TCPG | 14 | 26 | 4 |
| TCPD | 3 | 10 | 5 |
| AT2A1 | 2.5 | 41 | 5 |
| ENV1 | 5.5 | 32 | 6 |
| ASPH | 8.5 | 63 | 9 |
| TCPZ | 7 | 8 | 9 |
| CALX | 147 | 212 | 33 |
| NRROS | 220 | 688 | 382 |
| HYOU1 | 45 | 58 | N/A |
| UGGT1 | 20.5 | 43 | N/A |
| DJB11 | 10 | 20 | N/A |
| EF1G | 9.5 | 2 | N/A |
| TGFB1 | 9 | 27 | N/A |
| LRRC47 | 8.5 | 6 | N/A |
| EFTU | 8 | 3 | N/A |
| ACON | 6.5 | 2 | N/A |
| GANAB | 5.5 | 3 | N/A |

**Figure 1.**
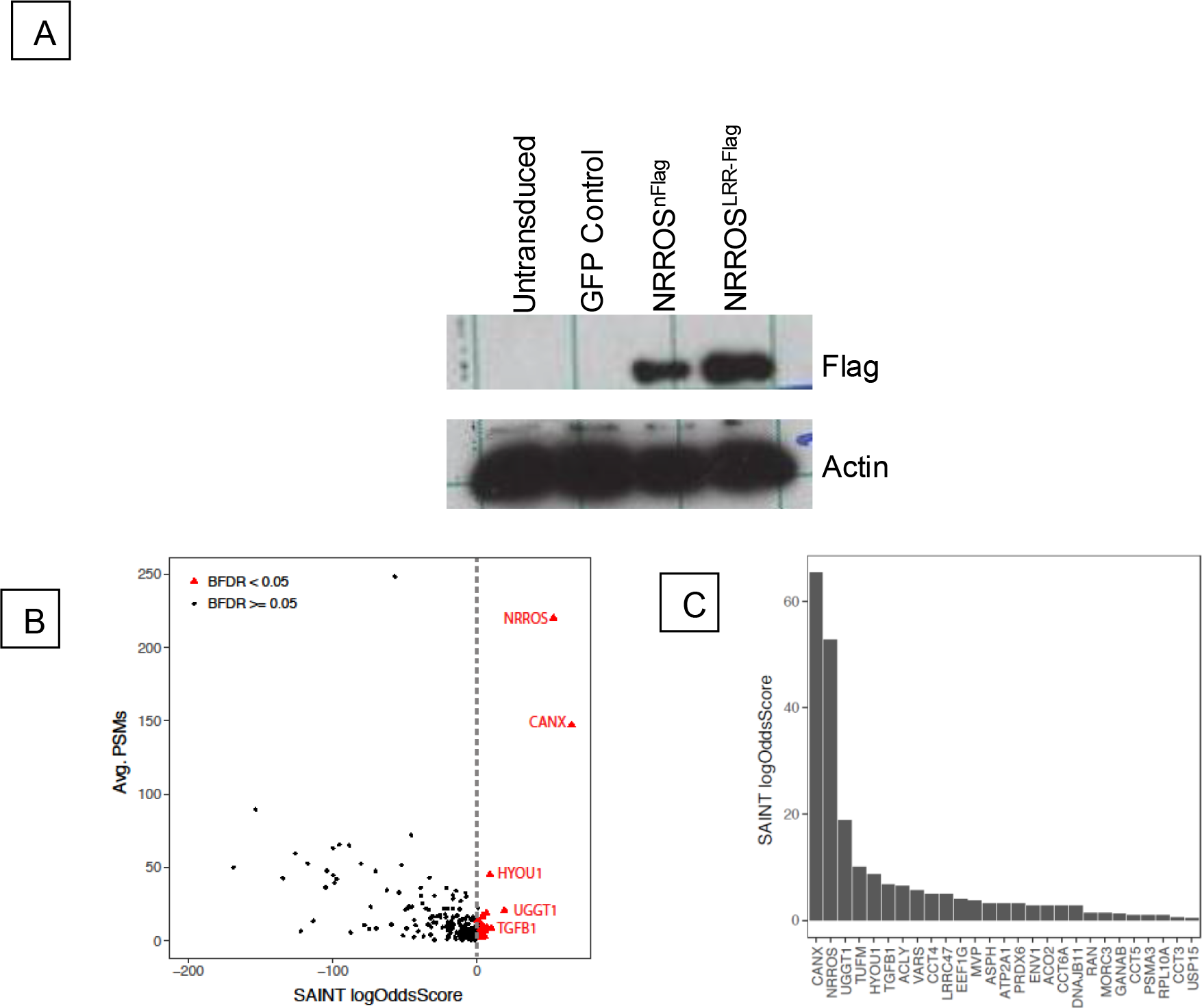
NRROS interactome in RAW264.7 cells. (A). Confirmation of NRROS^nFlag^ and NRROS^LRR-Flag^ by western blot analysis. (B). SAINT analysis of NRROS^Flag^ AP-MS data showing the logOddsScore (~specificity) vs. avg. PSMs (~abundance) for all identified proteins (n=2). Significant interactions (FDR < .05) are highlighted in red. Prominent abundant and significant interactions are labeled. (C). SAINT analysis of NRROS^Flag^ AP-MS data showing the logOddsScore for all significant proteins (FDR < .05).

**Figure 2.**
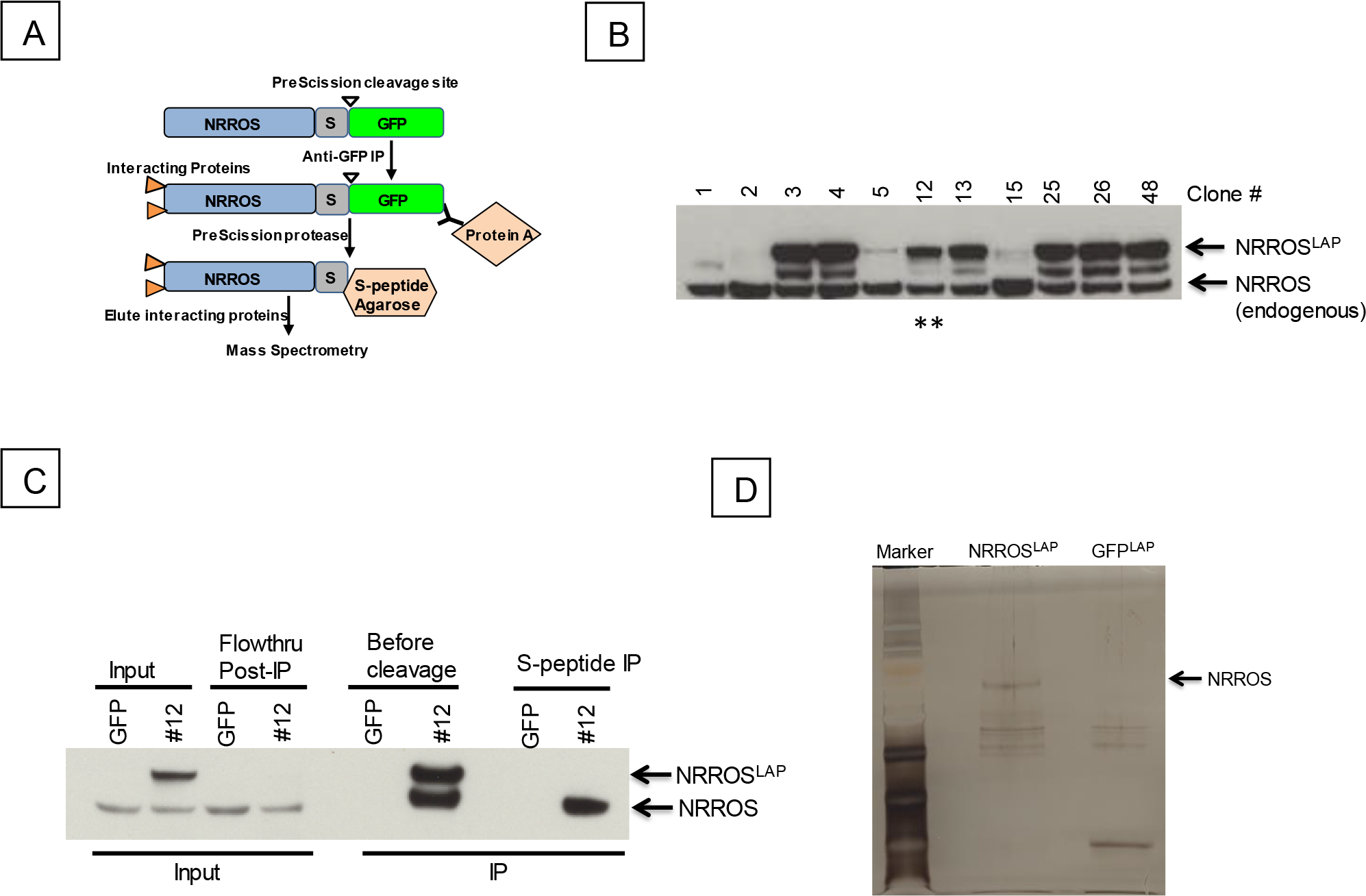
Generation and characterization of NRROS^LAP^ RAW264.7 cells. (A). Schematic of the Localization and affinity purification (LAP) tandem affinity purification strategy. S indicates S-peptide tag. (B). Screening of NRROS^LAP^ RAW264.7 single cell colonies for exogenous NRROS^LAP^ and endogenous NRROS by western blot analysis using anti-NRROS antibody. Clone 12 which was used for further analysis is marked with asterisk. (C). Confirmation of efficient NRROS^LAP^ immunoprecipitation by western blot analysis. (D). Visualization of NRROS^LAP^ enrichment by silver staining. NRROS is indicated, based on the molecular size observed in western blot analysis. Marker = Seablue Plus 2.

Next, to confirm our top hits from the mass spectrometric data, we immunoprecipitated NRROS^LAP^ from RAW cells and analyzed the interacting proteins by western blot. Consistent with the AP-MS data, calnexin was co-immunoprecipitated from NRROS^LAP^-expressing cells but not from control cells confirming a direct interaction between calnexin and NRROS (**Figure 4A**). We extended these studies to HEK293T cells where we overexpressed NRROS^nFlag^ and immunoprecipitated it using anti-Flag antibody. Again, consistent with previous results, several chaperone/co-chaperones such as calnexin, Bip and UGGT1 were co-immunoprecipitated with NRROS (**Figure 4B**). Additionally, we found NRROS to interact with Tgfβ1 when overexpressed together in 293T cells, confirming results from our AP-MS analyses and Bioplex 2.0 database (**Figure 4C**). We have previously reported that NRROS facilitates ERAD-mediated degradation of NOX2 monomers.^1^ When NOX2 and NRROS were co-overexpressed in HEK293T cells, we observed an increased interaction between NRROS and calnexin (**Figure 4D**). More importantly, we observed NRROS interaction with the mannosidase EDEM1 (ER degradation-enhancing alpha-mannosidase-like protein 1), a protein with known functions in the ERAD pathway,^18–21^ only under the conditions of increased NOX2 degradation (**Figure 4D**). Taken together, our data demonstrates that NRROS interacts with molecular chaperones such as calnexin and other ERAD-mediators and suggests that NRROS is involved in the regulation of protein biosynthesis and ER quality control (ERQC).

**Figure 3.**
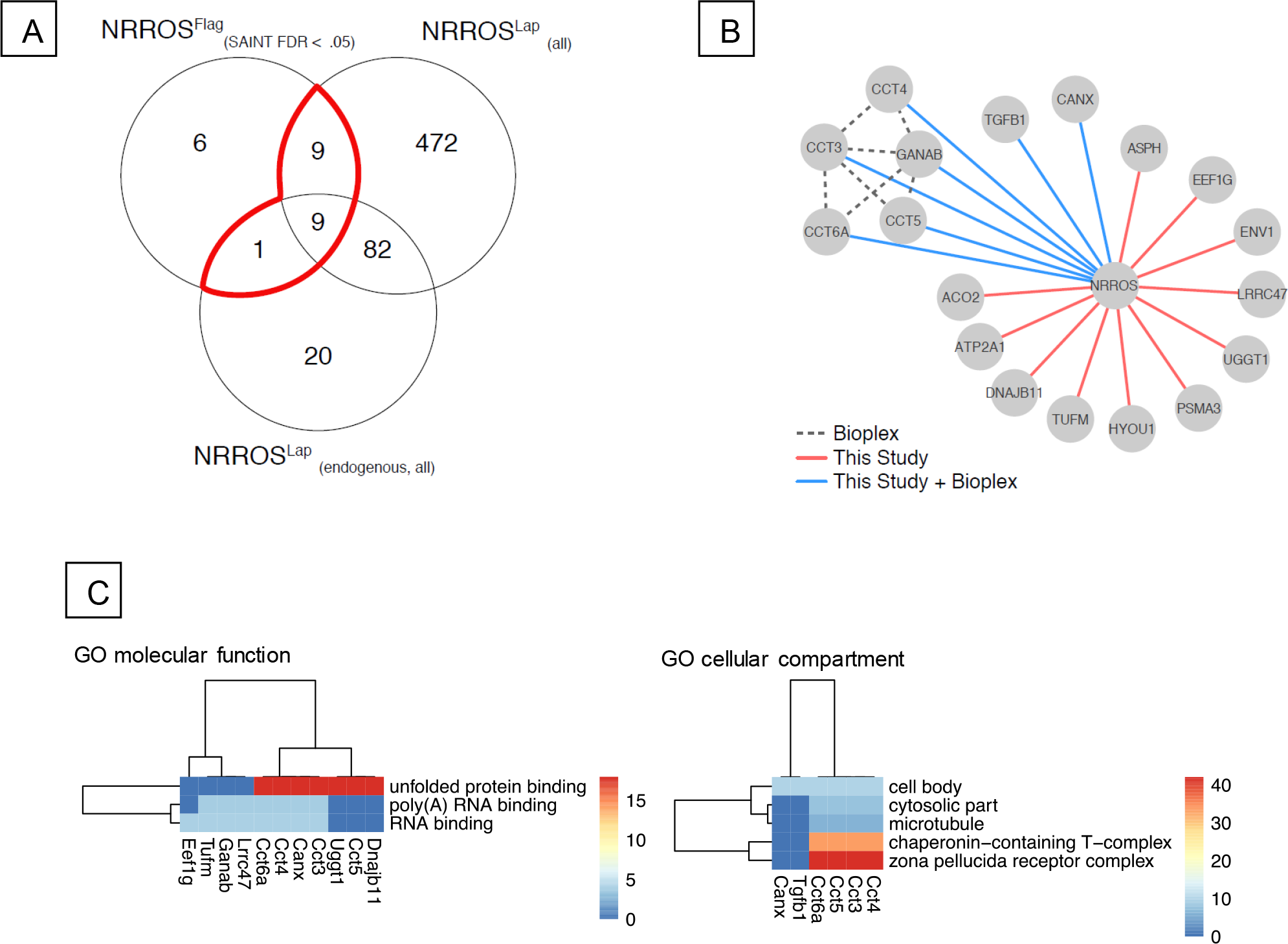
NRROS interactome in RAW264.7 cells detected using multiple approaches. (A). Overlap between significant interactions in NRROS^Flag^ (n=2, FDR < .05), NRROS^LAP^ over-expression (n=1, all proteins) and NRROS^LAP^ clone 12 (expression at endogenous level) (n=1, all proteins). The highlighted area encompasses all significant proteins in the NRROS^Flag^ AP-MS experiment that were identified in one or both NRROS^LAP^ AP-MS experiments. (B). The intersection obtained in (A) was integrated with the Bioplex 2.0 dataset and visualized as a protein interaction network. (C). Gene ontology enrichment analysis on the subset of significant interactions described in A. Significantly enriched annotations were filtered by p < 0.05, annotations with < 200 entries in GO and over-represented terms with >= 2 representations in the term. Since many genes matched multiple terms and vice versa, the Gena X Term matrix was clustered by the enrichment logOddsRatio.

**Figure 4.**
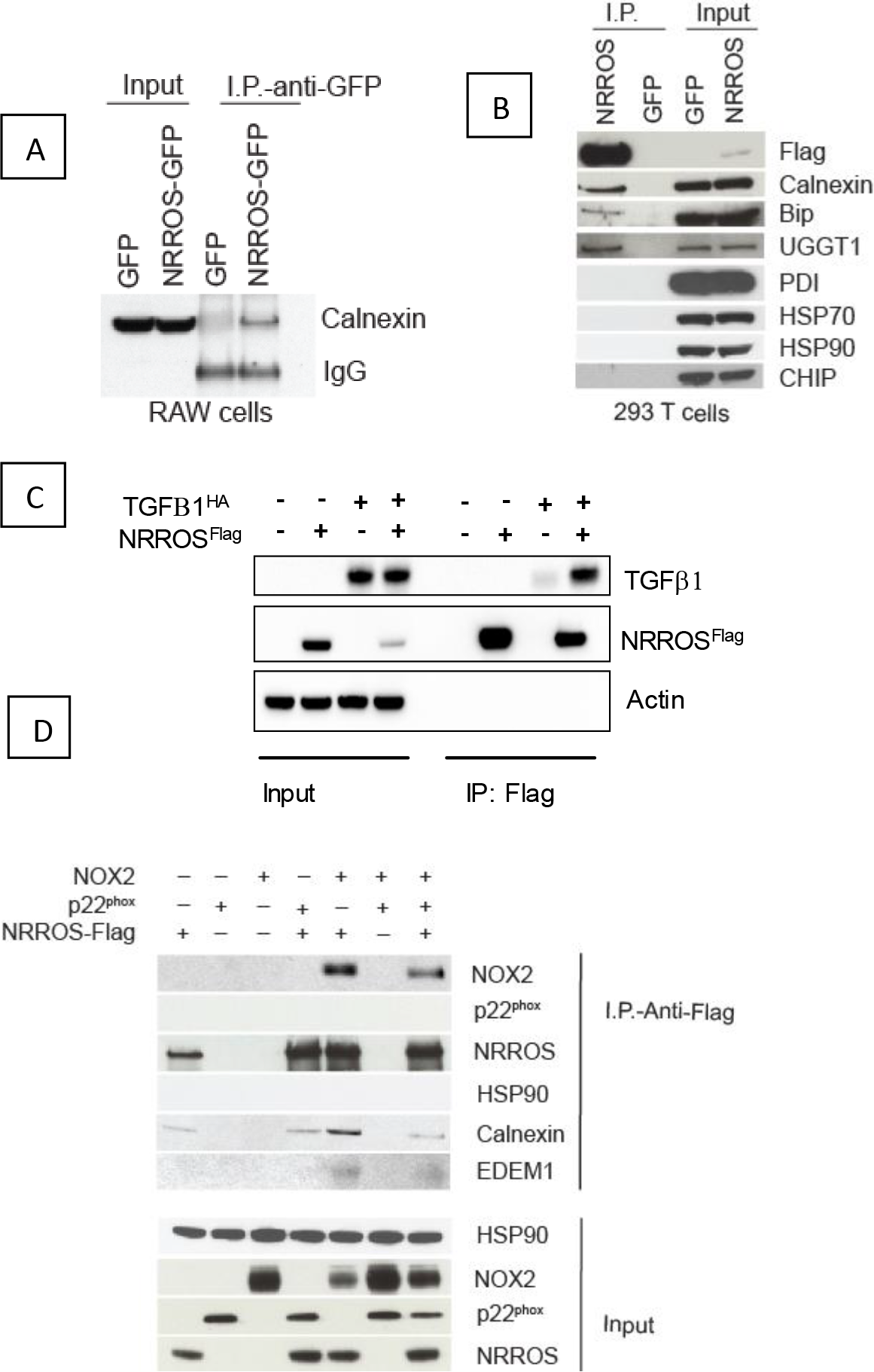
Confirmation of the NRROS interactome hits. (A). NRROS^LAP^ was immunoprecipitated from RAW264.7 cells using anti-GFP antibody and co-immunoprecipitation of endogenous calnexin was analyzed by western blot. (B). Co-immunoprecipitation calnexin, Bip, UGGT1 by NRROS^FLAG^ when overexpressed in HEK293T cells. Cell lysates were subjected to anti-Flag IP and analyzed by western blot analysis. No interaction was observed with PDI, HSP90, HSP70 or CHIP. (C). Co-immunoprecipitation of TGFβ1^HA^ by NRROS^FLAG^ when overexpressed in 293T cells. Cell lysates were subjected to anti-Flag IP and analyzed by western blot analysis. Actin is shown as a loading control (D). Co-immunoprecipitation of NOX2 and EDEM1 by NRROS^Flag^ when overexpressed in 293T cells. HSP90 is used as control.

## Discussion

In this report, we unravel NRROS interactome and show direct interactions between NRROS and molecular chaperones and ERAD mediators such as calnexin, Bip, UGGT1 and EDEM1. We employed multiple, complementary methods of co-immunoprecipitation and biochemical assays to profile a reproducible NRROS-interactome. Since NRROS is localized to the ER,^1^ it is conceivable that its interactome includes other ER-resident proteins. Moreover, comparison of NRROS interactome with that from BioPlex 2.0 database^17^ yields common interactors, providing an independent confirmation of our data. The GO term analysis of BioPlex 2.0 dataset indicated ER membrane/lumen and glycoprotein binding as the primary GO network for NRROS, similar to what we observed in our studies. An interesting interactor of NRROS, in addition to calnexin and Bip, common between the two studies is Hsp90b1/Cnpy3 (protein canopy homolog 3) complex. Hsp90b1 is a paralogue of HSP90.^22^ The Hsp90b1/Cnpy3 complex plays a critical role in proper folding and ER exit of multiple toll-like receptors^23^. This has important implications for NRROS in the regulation of innate immune responses. Another noteworthy NRROS-interactor common between our study and Bioplex 2.0 is TGFβ1. While we confirmed a direct interaction between NRROS and TGFβ1 when overexpressed together in HEK293T cells and in two of the three mass spectrometry analyses, we didn’t observe the interaction between NRROS and TGFβ1 in macrophages when NRROS was expressed at endogenous levels. It has been reported that NRROS expression is regulated by inflammatory signals.^1^ Therefore, tt is possible that the interaction between NRROS and TGFβ1 depends on NRROS expression levels and other physiological cues, which warrants further exploration. While we observed several proteins that are common in the NRROS-interactome between our study and Bioplex 2.0 database, there are few differences in the top hits. Some of the differences are attributable to different cell types utilized by the two studies to investigate the interactome. Our study investigated the NRROS interactome in macrophage cells that endogenously express the protein and hence, we believe, is physiologically relevant than an exogenous cellular expression system such as HEK293 cells that do not express *Nrros*.

We had previously demonstrated that NRROS recognizes un-dimerized NOX2 monomers and facilitates their degradation in a proteasome-dependent fashion.^1^ The NRROS interactors that we describe in the current study are proteins known to play important roles in the initial recognition of the ERAD substrates. Calnexin, for example, has been shown to interact with and retain incompletely assembled components of multi-subunit proteins, which eventually become substrates for ERAD.^24–26^ In this regard, we not only observed increased association of calnexin with NRROS under conditions that lead to NOX2 degradation in HEK293T cells but also co-immunoprecipitated EDEM1 with NRROS under these conditions (**Figure 4D**). EDEM1 has been shown to extract mis-folded glycoproteins from the calnexin cycle and facilitate their degradation through ERAD-mediated pathway.^18–19^ NOX2 dimerizes with p22^phox^ in phagocytic cells and this dimerization is essential for its stability.^12^ We also observed that NRROS interacts with NOX2 monomers which is a glycosylated protein but not with p22^phox^, a non-glycosylated protein^27 1^ (**Figure 4D**), suggesting that NRROS might play a role in recognizing some form of glycosylation signature on the proteins and enhance their degradation through ERAD. Furthermore, NRROS-deficient cells express higher NOX2 protein.^1^ Therefore, it is highly likely that, in NRROS-deficient cells, NOX2 undergoes repeated rounds of association with calnexin, thus increasing its chances of protein stabilization through p22^phox^ heterodimerization and hence increasing the protein expression whereas in NRROS-expressing cells, it interacts with NOX2 monomers and mediates their degradation. These data, taken together, suggest that NRROS is a molecular chaperone, with preferential expression in phagocytes, in the biosynthesis of multi-subunit proteins such as NOX2, either by itself or in association with calnexin. The functional importance of the NRROS interactome, particularly its interaction with calnexin and Bip is being investigated. We cannot rule out the possibility that some of the proteins present in NRROS-interactome are a result of NRROS itself undergoing glycosylation, since we have observed glycosylated NRROS (data not shown).

The ER is a specialized membrane bound organelle, with extensive network for the production of proteins in a cell.^28–31^ Proteastasis, the state of healthy, balanced proteome is dependent on the ER quality control (ERQC) machinery.^28^ ERAD, in addition to Unfolded Protein Response (UPR), is the primary ERQC system.^21, 32–35^ UPR and ERAD are vital to the health of a cell as evidenced by a large number of diseases linked to this pathway.^29, 36^ Our data suggests that NRROS plays an important role in the quality control of the ER, particularly that of multi-subunit proteins. More importantly, our data provides insights into the possibility of a broader set of proteins than the hitherto described NOX2 to be substrates for NRROS-mediated regulation through ERAD pathway and ERQC machinery.

## Acknowledgement

We thank Saikat Mukhopadhyay and Peter Jackson at Genentech for providing technical help and sharing reagents for LAP tandem affinity purification technique.

